# Colonization and extinction dynamics and their link to the distribution of European trees at continental scale

**DOI:** 10.1101/748483

**Authors:** Arnaud Guyennon, Björn Reineking, Jonas Dahlgren, Aleksi Lehtonen, Sophia Ratcliffe, Paloma Ruiz-Benito, Miguel A. Zavala, Georges Kunstler

## Abstract

**Aim:** Processes driving current tree species distribution are still largely debated. Attempts to relate species distribution and population demography metrics have shown mixed results. In this context, we would like to test the hypotheses that the metapopulation processes of colonization and extinction are linked to species distribution models.

**Location:** Europe: Spain, France, Germany, Finland, and Sweden.

**Taxon:** Angiosperms and Gymnosperms.

**Methods:** For the 17 tree species analyzed we fitted species distribution model (SDM) relating environmental variables to presence absence data across Europe. Then using independent data from national forest inventories across Europe we tested whether colonization and extinction probabilities are related to occurrence probability estimated by the SDMs. Finally, we tested how colonization and extinction respectively drive probability of presence at the metapopulation equilibrium.

**Results:** We found that for most species at least one process (colonization/extinction) is related to the occurrence probability, but rarely both.

**Main conclusions:** Our study supports the view that metapopulation dynamics are partly related to SDM occurrence probability through one of the metapopulation probabilities. However these links are relatively weak and the metapopulation models tend to overestimate the occurrence probability. Our results call for caution in model extrapolating SDM models to metapopulation dynamics.

## 1 Introduction

The vast majority of species have restricted geographical ranges (Holt & Keitt, 2000). Understanding the factors determining these ranges is fundamental to have insights on future species redistribution in the face of climate change. Species distribution is thought to be tightly connected to its ability to cope with local abiotic conditions and thus species’ niche (Pulliam, 2000; Thuiller, Lavorel, & Araújo, 2005; Soberón, 2007). This view underpins most statistical species distribution models (SDM) that relate species presence with local environmental conditions. These models have been extensively used in recent years and provided very detailed descriptions of species environmental requirements based on occurrence data. They provide, however, very little indication on how species distribution arises from population dynamics. This is surprising because we start to have a rich theoretical understanding of how population dynamics control species distribution and species range limits (Holt, Keitt, Lewis, Maurer, & Taper, 2005).

There are numerous routes through which range limits can arise (Holt et al., 2005). The first class of mechanisms consider only the local population dynamics when there is little effect of dispersal. The most classical view of this approach is that species ranges match the environmental conditions where birth rates exceed mortality rates (i.e. where the rate of population growth is above 1, Brown (1984)). Most existing field studies did not support this assumption, see the review by Pironon et al. (2017). For instance, Thuiller et al. (2014) demonstrated that major demographic parameters of European tree species were not strongly correlated to the occurrence probability derived from SDMs. Generally, only transplant experiments beyond species range have shown a tendency of a decrease in population growth rate or some demographic rates (Hargreaves, Samis, & Eckert, 2013; Lee-Yaw et al., 2016).

Then Holt et al. (2005) proposed two other classes of mechanisms based on local population dynamics: demographic stochasticity and temporal variability. Demographic stochasticity could increase the risk of extinction at the range limits (Boyce, Haridas, Lee, & Group, 2006; Ovaskainen & Meerson, 2010). For instance, this could be due to a lower absolute density leading to an increase in risk of extinction solely due to stochastic variability. However, several studies did not find strong support for the abundance center hypothesis which propose that abundance should be higher in the center of the distribution (Murphy, VanDerWal, & Lovett-Doust, 2006; Sagarin, Gaines, & Gaylord, 2006). Temporal variability could also increase the risk of local extinction even if the average growth rate and the average population size are not limiting factors. Rare extreme conditions or highly unstable environmental conditions might control the extinction risk at the range limits. Field tests of these mechanisms are extremely rare, and show weak support for this hypothesis. Csergő et al. (2017), using detailed demographic data for plant species (including trees), found no clear link between climate suitability, derived from a SDM, and several detailed population metrics including time to quasi-extinction, stochastic population growth rate or transient population dynamics.

Another class of mechanisms underlying species ranges is a regional equilibrium between colonization/extinction dynamics of populations connected by dispersal (Holt et al., 2005; Holt & Keitt, 2000). This last class relates to the metapopulation paradigm and proposes that species ranges arise from the gradient of three variables: the extinction rate, the colonization rate, and the habitat structure (i.e. the availability of suitable area for settlement). In this model, the dynamic of the local population is ignored when compared to regional dynamics (Drechsler & Wissel, 19997). Such models thus ignore the details of the population dynamics but rather focus on patch occupancy dynamics (extinction and colonization events). Few studies have focused on these processes for tree species (see Purves, Zavala, Ogle, Prieto, & Benayas, 2007; García-Valdes, Zavala, Araújo, & Purves, 2013; García-Valdés, Gotelli, Zavala, Purves, & Araújo, 2015; Talluto, Boulangeat, Vissault, Thuiller, & Gravel, 2017). Results in North America (Talluto et al., 2017) showed that metapopulation processes captured potential future range shifts for most tree species. García-Valdes et al. (2013) inferred potential changes in species distribution in Spain, but we still lack studies that explore this mechanism at the European scale, covering a larger portion of different species distribution.

Here we propose to analyze how local species extinction and colonization probabilities vary within the range of the the main European tree species across the entire continent using more than 80 000 plots of national forest inventories. Species distributions are summarized by occurrence probability estimated with an ensemble SDM fitted to independent data extracted from the EU-Forest data set (Mauri, Strona, & San-Miguel-Ayanz, 2017). We use presence/absence from NFI data to get observations of extinction and colonization events, a colonization event being then separated into an outcome of a seed input and a successful recruitment.

We then analyze the relationship between the occurrence probability derived from SDMs and the extinction/recruitment probabilities derived from NFI data to test the following hypotheses:

- Extinction probabilities increase when the SDM occurrence probability decreases.
- Recruitment probabilities decrease when the SDM occurrence probability decreases.
- Finally we analyse how the equilibrium occurrence probability, predicted by metapopulation models using estimates of extinction/recruitment probability, match the current SDM occurrence probability. This allows us to evaluate the relative importance of extinction and recruitment in driving the distribution of each species.

## 2 Materials and Methods

Our objective is first to test how extinction and recruitment probabilities vary as a function of the SDM derived occurrence probability for the dominant European tree species. Then we derived a potential equilibrium and compared it to current SDM occurrence probability to analyse the relative importance of each of the two processes, and whether it under or overestimates the current occurrence probability.

To do this, we first gathered data on tree local extinction and colonization events from national forest inventory plots. Then, we estimated the occurrence probability with SDM models fitted to independent data extracted from the EU-Forest database. Subsequently, we modelled extinction and recruitment probabilities in function of SDM occurrence probability with two observations of occupancy data via a spatially inhomogeneous Markov chain. Because national forest inventories provide little information on the local seed source around the plots, we used estimation of species local frequency in 1 km grid directly from JRC maps (see section 2.2) as a surrogate of seed source. Finally, we derived the probability of presence at equilibrium based on the estimated extinction and recruitment considering two alternative formulations: one in which communities are considered as closed systems, the other in which communities are open to external seed sources.

### 2.1 NFI datasets

To calibrate our model, we required information on the presence/absence of each species at two different dates over a large geographical area to cover, as far as possible, the entire species distributions. We used a database of tree data from the National Forest Inventories (NFI) of Finland, France, Germany, Spain, Sweden, compiled as part of the FunDivEurope project (http://www.fundiveurope.eu, Baeten et al. (2013)).

Inventory protocols differs between NFIs (see Supplementary materials Section 1 for a detailed description of each survey protocol). Surveys were conducted within a circular plot with a fixed radius or in concentric subplots with different radius and minimum diameter at breast height (DBH) for all NFIs except Germany, where an angle-count method (basal area factor of 4 *m*^2^*ha*^−1^) was used. Because the DBH thresholds for trees to be recorded varied between the inventories, we only included trees with a DBH of 10 cm or greater. For each NFI, except France, two inventory surveys were conducted with a variable time interval (from 4 to 16 years, see Figure 1 b). The French inventory is based on single surveys where the growth of alive trees (based on a short core) and approximate the date of death of dead trees are estimated and can be used to reconstruct the stand structure five years before the census, making it comparable with revisited plots data.

**Figure 1:**
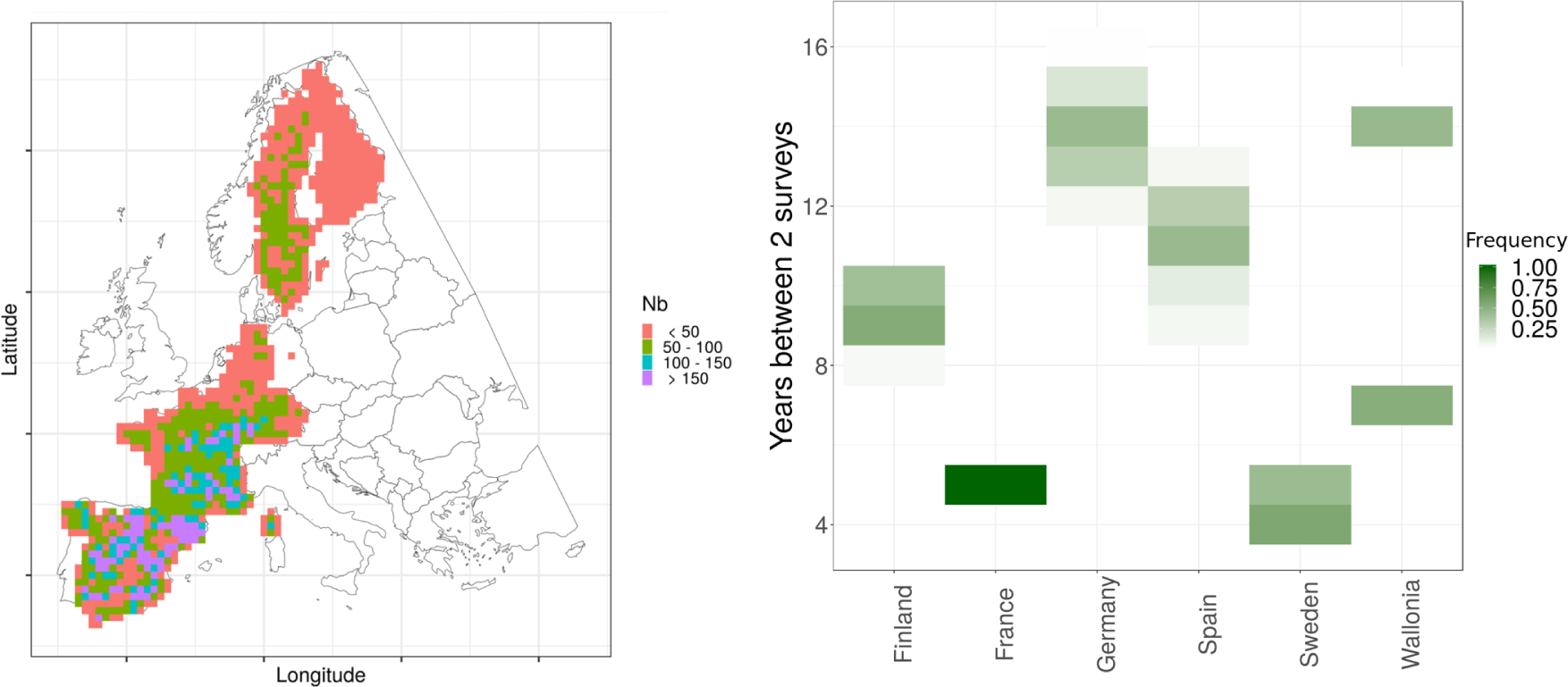
(a) Density map of NFI plots (grid of 50 km x 50 km) used to estimate colonization and extinction events. (b) Distribution of number of years between subsequent surveys by country

To avoid influences of management on the extinction and colonization events, we discarded plots where a management effect was reported between surveys. This led to a selection of 80 157 plots with 173 species. Among these species we selected the most abundant species (the cumulative basal area of species retained represented more than 95% of the total basal area) and excluded exotic species as well as species for which JRC maps (see below) were not available.

For each plot, a species was considered present when at least one tree was observed. The succession of two surveys allowed then to deduce state transitions (0 → 1 for local colonization, 1 → 0 for local extinction). Since several protocols are based on concentric circular plots with varying DBH thresholds, a newly observed tree might not be a recruited tree, i.e. its DBH during the first census was above 10 cm, but it was not recorded due to the larger DBH threshold for its subplot. We used a species specific growth model to estimate the probability that a new tree (present only in the second census) had a former DBH below 10 cm. The growth model was built as a generalized linear model using an aridity index, the sum growing degree days, and tree DBH as explanatory variables (see Supplementary Materials Section 2). We thus considered a plot as colonized if the probability that the largest newly observed tree had a former DBH below 10 cm greater or equal to 0.5, otherwise the species was considered as present at both censuses (1 → 1). This correction had a strong impact on the Spanish and German inventories, significantly reducing the number of colonization events. We decided to exclude from further analysis species with less than 10 events for extinction or colonization (*i.e. Quercus suber, Pinus pinea* and *Acer pseudoplatanus*), resulting in a final selection of 17 species.

### 2.2 Joint research center - species local frequency

The density of NFI plots is too low to accurately describe the local abundance of trees that can disperse seeds into a given plot. Distances between NFI plots are about 1 km or above whereas most dispersal events occur in less than 100 m from the seed source (see for example Nathan, Safriel, and Imanuel (2001), Bullock et al. (2017)). To represent seed inputs into a plot, we thus used the species’ local frequency (hereafter *F*_*JRC*_) in the corresponding 1 km cell produced by the Joint Research Center (RPP - Relative Probability of Presence on JRC website (https://forest.jrc.ec.europa.eu/en/european-atlas/atlas-data-and-metadata/), see San-Miguel-Ayanz, de Rigo, Caudullo, Houston Durrant, and Mauri (2016). Each map estimates the relative frequency of the species based on datasets of field observations as represented in the Forest Information System for Europe (FISE), which integrates National Forest Inventories, BioSoil and Forest Focus data sets. The presence/absence data are assimilated at a spatial resolution of 1 km based on multiple smoothing kernels of varying dimension. Independent estimations of forest cover extracted from the Pan-European Forest Type Map 2006 (FTYP2006, http://forest.jrc.ec.europa.eu/forest-mapping/forest-type-map) are used to rescale the species frequency by the cover of broadleaved forest, coniferous forest or other non-forest categories based on 25 m x 25 m pixels (San-Miguel-Ayanz et al., 2016). We chose this variable because it summarized a very large amount of data on a European scale and can be considered as a strong indicator of the proportion of adjacent plots in which the species is present within a 1 km patch. An explicit representation of seed availability via dispersal mechanisms was beyond the scope of this work because available data do not provide a detailed description of the seed source in the plot surroundings from where most dispersal events occur (Nathan et al., 2001). However, because the JRC species local frequency data is based on the spatial integration of presence-absence observations, our seed source estimate can be influenced by data beyond the 1 km grid. The long-distance seed dispersal events are thus not excluded, even if there is no observation of presence in the 1 km cell.

### 2.3 SDM

We estimated species occurrence probability (hereafter *P*_*occ*_) on each NFI plot with ensemble species distribution models fitted to the EU-Forest data set (Mauri et al., 2017) which provides species presence/absence on a 1 km grid. The initial EU-Forest data set includes more than 250 000 plots across Europe including countries not present in FUNDIV. We excluded all NFI observations from the EU-Forest to avoid using the same data to estimate both extinction/colonization and the SDM probability of presence. This exclusion was performed to avoid any potential circularity arising from the use of the same data in both analyses. After excluding NFI observations, we retained 9600 data points across Europe, coming from ForestFocus and BioSoil campaigns. For each grid point, we extracted mean annual temperature, precipitation of wettest quarter, temperature and precipitation seasonality from CHELSA climatologies (Karger et al., 2017), pH measured in water solution (5 cm depth) from SoilGrid (Hengl et al., 2017), and aridity index (the mean annual precipitation divided by the mean annual potential evapotranspiration) and actual evapo-transpiration from CGIAR-CSI (Trabucco & Zomer, 2010). We verified that correlation coefficients between variables were always lower than 0.7 (Dormann et al., 2013), except for pH and mean annual temperature which had a correlation coefficient of 0.72. We nevertheless decided to keep both variables, as the correlation was only slightly higher than 0.7, and both soil information and mean temperature variables may be important drivers for species distribution. Then we fitted ensemble SDM models with BIOMOD2 (Thuiller, Lafourcade, Engler, & Araújo, 2009) using four different models (GAM, GLM, GBM, and Random Forest). Based on this ensemble model we estimated species occurrence probability on each NFI plot for all species. Details on the evaluation on the predictive power of the SDM are provided in the Supplementary materials Section 3 (see Figure 1 with performance scores of SDM for each species based on True Skill Statistic, TSS and Area Under the Curve of the Receiver Operating Characteristic, AUC).

### 2.4 Patch occupancy transition model

The patch occupancy model is a spatially inhomogeneous Markov chain, the state vector being the patch occupancy of the N plots *X*(*t*) at time t. The probability of transition between the two time successive patch occupancy patterns is:

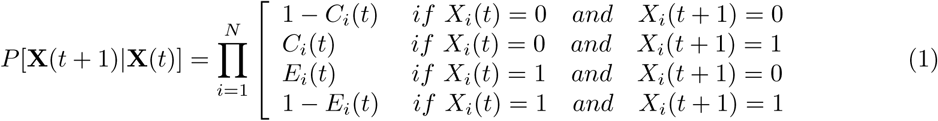

where N is the total number of plots observed, *E*_*i*_ the extinction probability in plot *i*, and *C*_*i*_ the colonization probability in plot *i*.

The extinction probability (E) of a species in a plot only depends on the local environmental conditions, i.e. the occurrence probability derived from the SDM (*P*_*occ*_). The colonization probability (C) is divided into two contributions: recruitment probability (R) which depends on *P*_*occ*_, and seed source (S). The recruitment probability R is the probability of at least one tree reaching 10 cm between two protocols. Colonization probability is simply expressed as the product of *R* and *S*, where the seed source *S* is estimated by the JRC as presented above. Colonization events can occur in any plot with a non-null seed source.

Recruitment (R) and extinction (E) probabilities were related to the SDM occurrence probability *P*_*occ*_ and the species local frequency *F*_*JRC*_ as follows:

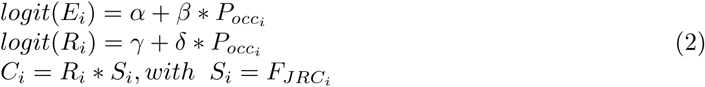

Differences in protocols between countries can influence the probability of observing extinction and colonization events. To account for this protocol effect in our analysis we used fixed country specific intercept parameters (*α* and *γ*).

Because the time interval between two censuses may vary across plots (between 4 and 15 years), we standardized the parameters to a 5 years sampling interval as done in Talluto et al. (2017), the probability of an recruitment/extinction was computed as:

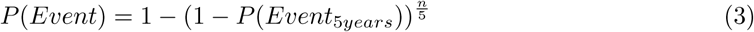

with n being the number of years between the two censuses.

### 2.5 Calibration of the model

For each species, extinction and recruitment parameters were estimated separately using a Metropolis Hastings Monte Carlo sampling algorithm, with priors following a Cauchy distribution (Gelman, Jakulin, Pittau, & Su, 2008; Ghosh, Li, & Mitra, 2018) using JAGS (Plummer, 2003). Convergence was checked by evaluating whether the Gelman-Rubin convergence statistic was below 1.1, as recommended by Brooks and Gelman (1998), using 4 chains.

### 2.6 Probability of presence at equilibrium

Finding a link between either recruitment or extinction and the SDM *P*_*occ*_ does not necessarily mean that the equilibrium model would yield the same occurrence probability as the SDM. To evaluate this, we derived from the estimates of recruitment and extinction a probability of presence at equilibrium (hereafter *P*_*eq*_). We explored the match with *P*_*occ*_ and the relative contribution of extinction and recruitment probabilities. The equilibrium can be defined in two ways: (1) We can assume that grid cells are open systems with a fixed seed source *S*, where the probability of presence in the grid cell is a function of extinction, recruitment, and the value of seed source. In this case there is no feedback of the colonization and extinction on the seed source. (2) We can assume that grid cells are closed systems of interconnected suitable patches, with a feedback of the colonization and extinction processes on the seed source. In this case, an extinction probability exceeding the colonization probability would lead to a species absence.

Both types of equilibrium can be derived from the same equation:

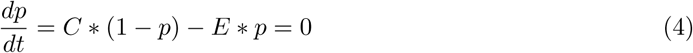

with p the proportion of suitable patches occupied.

The difference between the two types of equilibrium corresponds to different formulations of *C*. In the first formulation, *S* is constant over time and *C* = *R* × *S*, while in the second formulation, *S* varies with *P*_*occ*_ and *C* = *p* ∗ *R*. These two alternative formulations lead to the following equilibria:

- (1) when we consider a fixed seed source, and compute the equilibrium state for each plot: 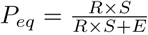
- (2) when we consider that the seed source is linked to the proportion of occupied patches within each 1 km grid cell, then the proportion of suitable occupied patches is 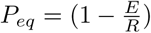.

For both formulations, we studied the relative impact of extinction and colonization (including seed source and recruitment probability) on the equilibrium state by fixing one of the probabilities to its mean and letting the other vary based on our estimated slope of response to the SDM occurrence probability. We also computed the probability at equilibrium, letting both extinction and colonization vary with *P*_*occ*_. In the first model we can also set the fixed seed source to one (no dispersal limitation) or let the fixed seed source vary with *P*_*occ*_ based on their observed links.

## 3 Results

### 3.1 Recruitment/Extinction dependencies

Results show that at least one of the estimated probabilities (recruitment or extinction) is significantly related to the SDM occurrence probability for all species, with the exception of *Fraxinus excelsior*.

Overall, recruitment probability increases with the SDM occurrence probability (*δ* is positive, Figure 2 left). The slope for the recruitment model is positive when considering all species posteriors, and all species but *Abies alba* have a positive mean slope value. However, the effect is significant (at the 5% level) for only nine species out of 17.

**Figure 2:**
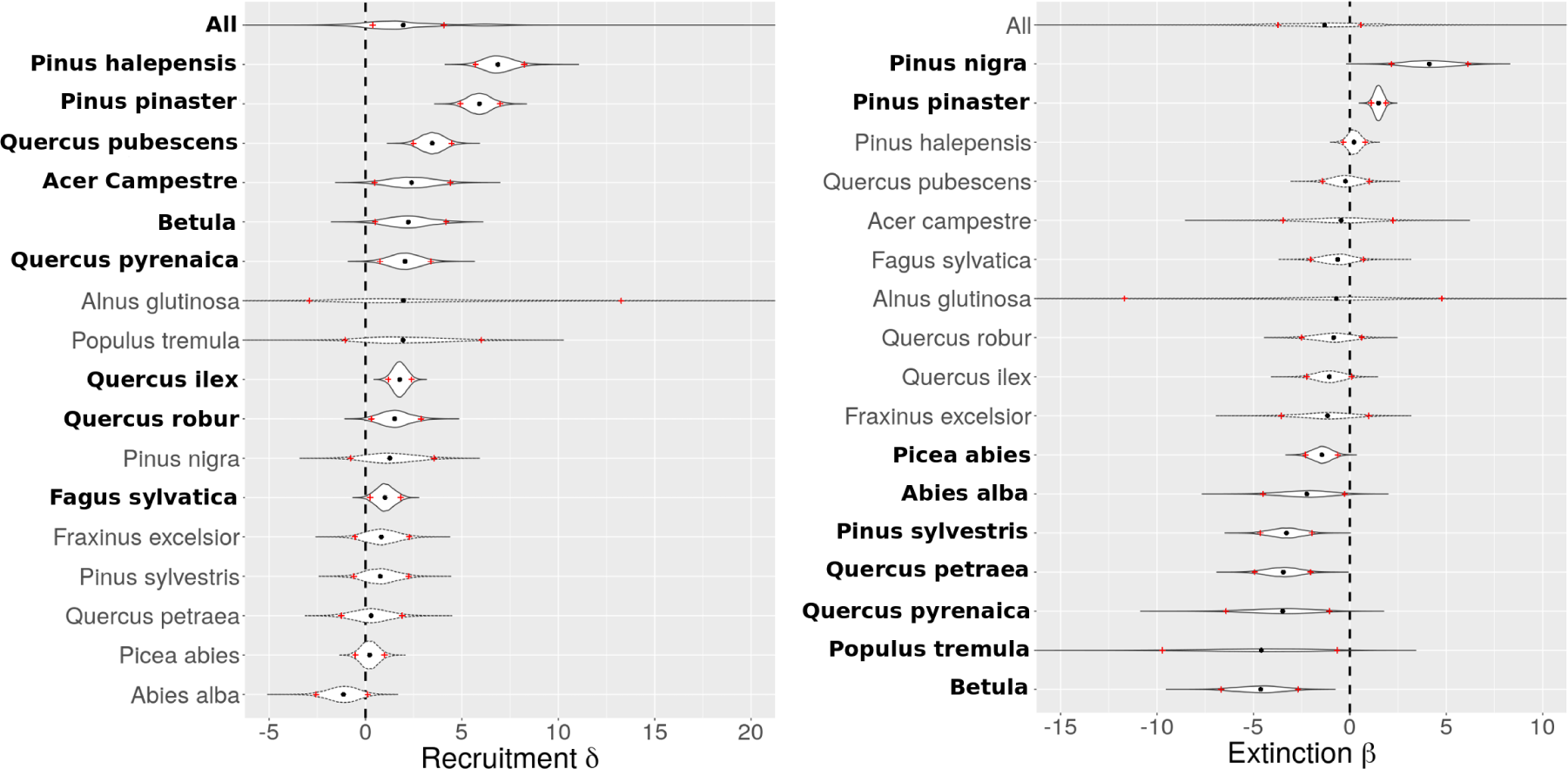
Posterior distribution of the slope of response of recruitment (left, *δ*) and extinction (right, *δ*) to *P*_*occ*_. Black points are posterior medians, red crosses indicate the 5th-95th percentile intervals. Species with name in bold have their interval not crossing 0.

Extinction probability is not significantly influenced by occurrence probability when considering all species. *Pinus nigra* and *Pinus halepensis* both present a positive slope, and seven species have a significant negative slope.

*Populus tremula* and *Alnus glutinosa* exhibits a very broad posterior for the slope parameters which can be related to the small range of probabilities of occurrence and the relative low discriminative power of their SDM (see Supplementary materials Section 4).

Model performances according to the True Skill Statistics (TSS, see (Allouche, Tsoar, & Kadmon, 2006)) varied from good (*TSS* > 0.5), average (0.3 < *TSS* < 0.5), to poor (*TSS* < 0.3) depending on the species and process (Tables 1 and 2).

**Table 1:** Estimates of *δ*, the slope of recruitment response to *P*_*occ*_ per species and their 90% confidence interval (see Materials and Methods for details on the model). ΔDIC is the difference of deviance information criterion – DIC – between the model and a null model (a model including only fixed country effects). TSS is the True Skill Statistics. Nb of events is the number of colonization events. TSS are calculated using the median of parameter posterior distributions.

| Species | Nb of events | Median $\delta$ | 90% Interval $\delta$ | TSS | $\Delta$ DIC |
| --- | --- | --- | --- | --- | --- |
| *Pinus sylvestris* | 149 | 0.8 | -0.6/2.2 | 0.61 | 0.3 |
| *Picea abies* | 231 | 0.2 | -0.5/1.0 | 0.70 | 1.7 |
| *Fagus sylvatica* | 160 | 1.0 | 0.2/1.8 | 0.52 | -2.9 |
| *Quercus robur* | 103 | 1.5 | 0.3/2.9 | 0.58 | -3.2 |
| *Quercus petraea* | 48 | 0.3 | -1.3/1.9 | 0.51 | 1.3 |
| *Pinus pinaster* | 66 | 5.9 | 4.9/7.0 | 0.76 | -107.5 |
| *Quercus ilex* | 351 | 1.8 | 1.2/2.4 | 0.58 | -23.0 |
| *Pinus nigra* | 48 | 1.3 | -0.8/3.6 | 0.67 | 0.2 |
| *Abies alba* | 53 | -1.1 | -2.6/0.10 | 0.60 | -0.5 |
| *Pinus halepensis* | 50 | 6.9 | 5.7/8.2 | 0.83 | -120.0 |
| *Quercus pubescens* | 87 | 3.5 | 2.5/4.5 | 0.73 | -35.0 |
| *Betula* | 265 | 1.3 | 0.5/4.2 | 0.63 | -2.9 |
| *Fraxinus excelsior* | 121 | 0.8 | -0.6/2.3 | 0.35 | 0.7 |
| *Quercus pyrenaica* | 85 | 2.0 | 0.8/3.4 | 0.79 | -6.1 |
| *Alnus glutinosa* | 47 | 2.0 | -2.9/13.3 | 0.54 | -0.2 |
| *Populus tremula* | 73 | 1.9 | -1.1/6.0 | 0.48 | -0.5 |
| *Acer campestre* | 97 | 2.4 | 0.5/4.4 | 0.67 | -2.9 |

**Table 2:**
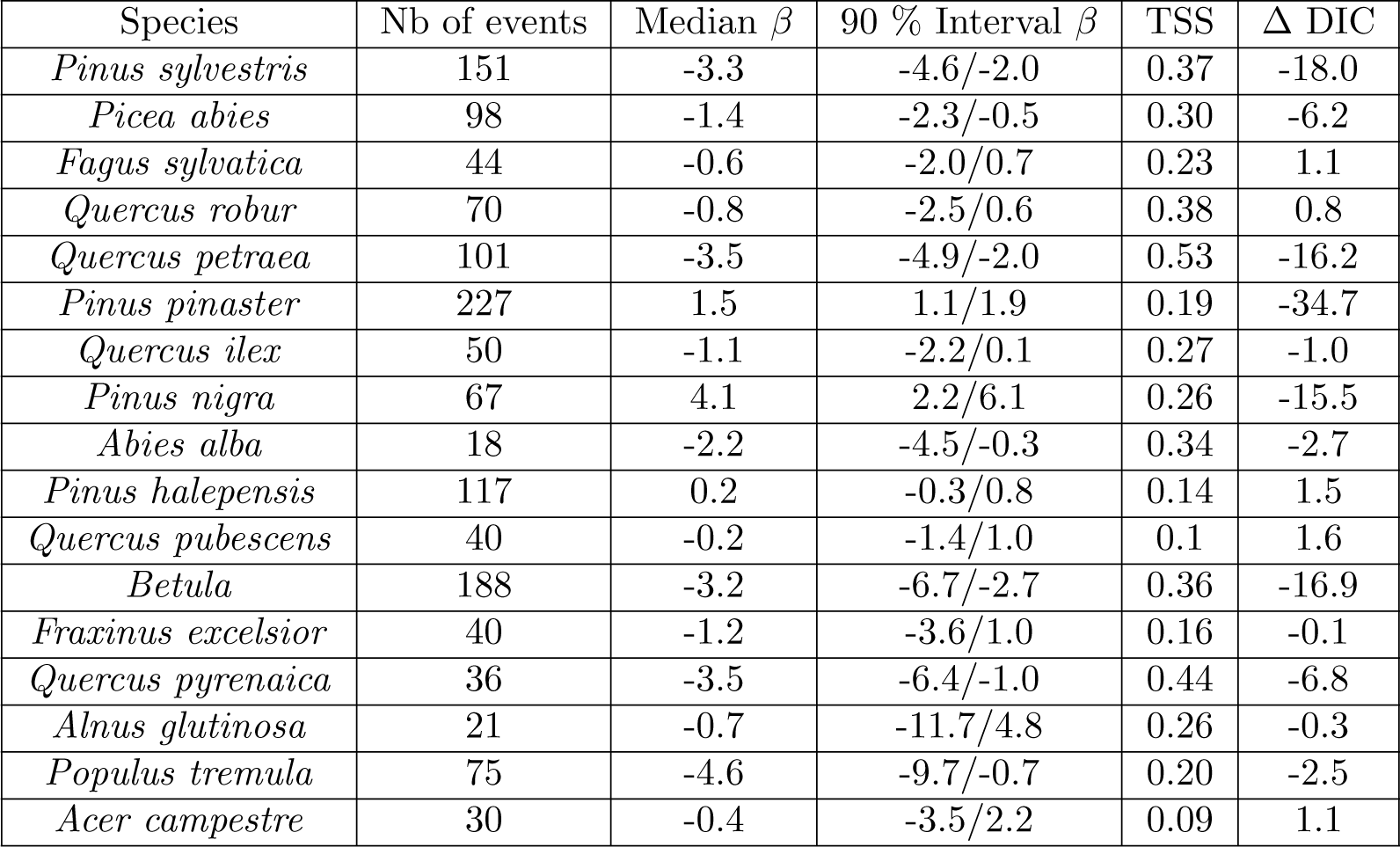
Estimates of *δ*, the slope of response of extinction to *P*_*occ*_ per species and their 90% confidence interval (see Materials and Methods for details on the model). ΔDIC is the difference of deviance information criterion – DIC – between the model and a null model (without *P*_*occ*_ dependency). TSS is the True Skill Statistics. Nb of events is the number of extinction events. TSS are calculated using the median of parameter posterior distributions.

| Species | Nb of events | Median $\beta$ | 90 % Interval $\beta$ | TSS | $\Delta$ DIC |
| --- | --- | --- | --- | --- | --- |
| *Pinus sylvestris* | 151 | -3.3 | -4.6/-2.0 | 0.37 | -18.0 |
| *Picea abies* | 98 | -1.4 | -2.3/-0.5 | 0.30 | -6.2 |
| *Fagus sylvatica* | 44 | -0.6 | -2.0/0.7 | 0.23 | 1.1 |
| *Quercus robur* | 70 | -0.8 | -2.5/0.6 | 0.38 | 0.8 |
| *Quercus petraea* | 101 | -3.5 | -4.9/-2.0 | 0.53 | -16.2 |
| *Pinus pinaster* | 227 | 1.5 | 1.1/1.9 | 0.19 | -34.7 |
| *Quercus ilex* | 50 | -1.1 | -2.2/0.1 | 0.27 | -1.0 |
| *Pinus nigra* | 67 | 4.1 | 2.2/6.1 | 0.26 | -15.5 |
| *Abies alba* | 18 | -2.2 | -4.5/-0.3 | 0.34 | -2.7 |
| *Pinus halepensis* | 117 | 0.2 | -0.3/0.8 | 0.14 | 1.5 |
| *Quercus pubescens* | 40 | -0.2 | -1.4/1.0 | 0.1 | 1.6 |
| *Betula* | 188 | -3.2 | -6.7/-2.7 | 0.36 | -16.9 |
| *Fraxinus excelsior* | 40 | -1.2 | -3.6/1.0 | 0.16 | -0.1 |
| *Quercus pyrenaica* | 36 | -3.5 | -6.4/-1.0 | 0.44 | -6.8 |
| *Alnus glutinosa* | 21 | -0.7 | -11.7/4.8 | 0.26 | -0.3 |
| *Populus tremula* | 75 | -4.6 | -9.7/-0.7 | 0.20 | -2.5 |
| *Acer campestre* | 30 | -0.4 | -3.5/2.2 | 0.09 | 1.1 |

Recruitment models showed average to good performance for all species. Extinction models showed average to good performance for seven (41 %) species. Model scores were not related to the number of observations (p-values of 0.69 and 0.48 for extinction and recruitment respectively). We also computed Δ*DIC* for each process and species as a complementary quantification of model performance (Spiegelhalter, Best, Carlin, & Van Der Linde, 2002). DIC helps compare the relative fit of models, and in our case a negative value support the inclusion of *P*_*occ*_ dependency in the model.

Given the scarcity of colonization or extinction events, we also tested the robustness of our slope estimates to the proportion of zeros by refitting the model after controling the proportion of zeros in the data (see Supplementary materials Section 5).

Since the range of *P*_*occ*_ is different between species, and the link function is non linear, the slope is not sufficient to evaluate the magnitude of recruitment and extinction variability. We thus also computed the relative contribution of *P*_*occ*_ to extinction and recruitment (Figure 3) as the difference between the probability of extinction (colonization) at the low *vs.* high end of *P*_*occ*_ (respectively 5 and 95 % percentiles).

**Figure 3:**
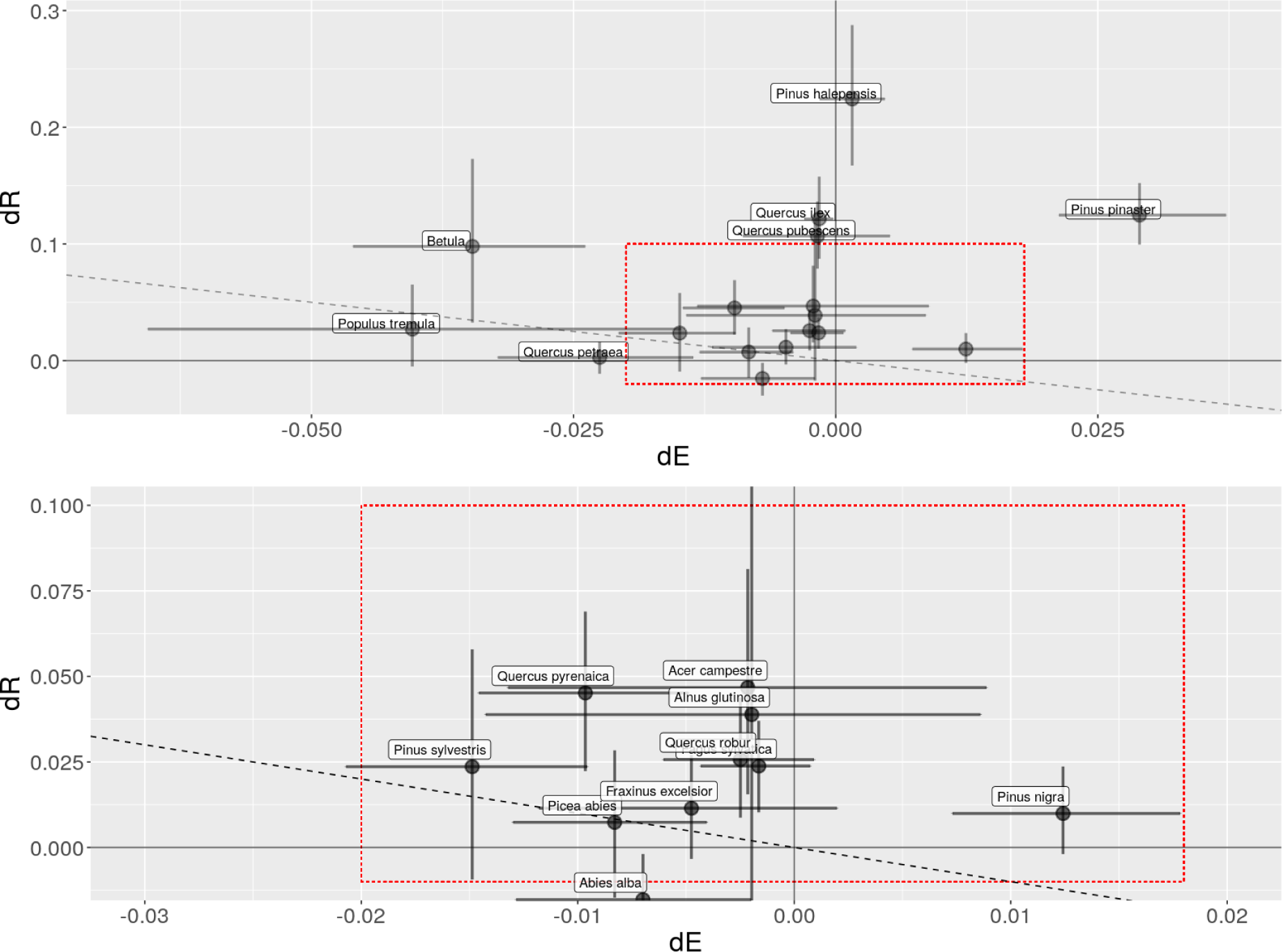
Relative contribution of *P*_*occ*_ to recruitment/extinction probabilities (dR and dE, respectively). For each species, dE and dR are calculated as the differences at high *P*_*occ*_ (95th centile) and low *P*_*occ*_ (5th centile). Bottom Figure is a zoom of the top Figure, indicated by a red square. Negative dE means a higher extinction rate at low occurrence probability; positive dR means a lower colonization at low occurrence probability. On both plots dashed line represents *dR* = −*dE*.

For most species, the relative contribution was higher for recruitment than for extinction, i.e. most species are above the diagonal in the Figure 3, particularly for *Quercus ilex, Quercus pubescens* and *Pinus halepensis*. Only *Quercus petraea* and *Abies alba* were below the diagonal, with a higher relative contribution of *P*_*occ*_ on the extinction than on the recruitment probability.

### 3.2 Equilibrium

The probabilities of both colonization and extinction depend on the SDM occurrence probability *P*_*occ*_. As a consequence, the probability of presence at equilibrium *P*_*eq*_ is directly a function of *P*_*occ*_. However, the shape of the function and the match between *P*_*eq*_ and *P*_*occ*_ depend on the estimates of the slopes and intercepts of the colonization and extinction models (see Supplementary materials section 6).

The relationship between *P*_*eq*_ and *P*_*occ*_ was positive for most species when we accounted for the variation of both recruitment and extinction probability (green curve in figure 4). Only *Pinus nigra* had a negative relationship. *P*_*eq*_ showed few variations and overall overestimated *P*_*occ*_. When dispersal limitation is not included, *P*_*eq*_ is above 0.5 when colonization probability exceeds extinction probability, which is always the case for all species. If we included the seed approximation in the formulation (black curves in figure 4), the match between *P*_*eq*_ and *P*_*occ*_ was stronger. Only *Quercus ilex* exhibited systematically higher *P*_*eq*_ than *P*_*occ*_, while for *Quercus petraea P*_*eq*_ tended to be lower than *P*_*occ*_. For all other species, *P*_*occ*_ stood within the range of *P*_*eq*_.

**Figure 4:**
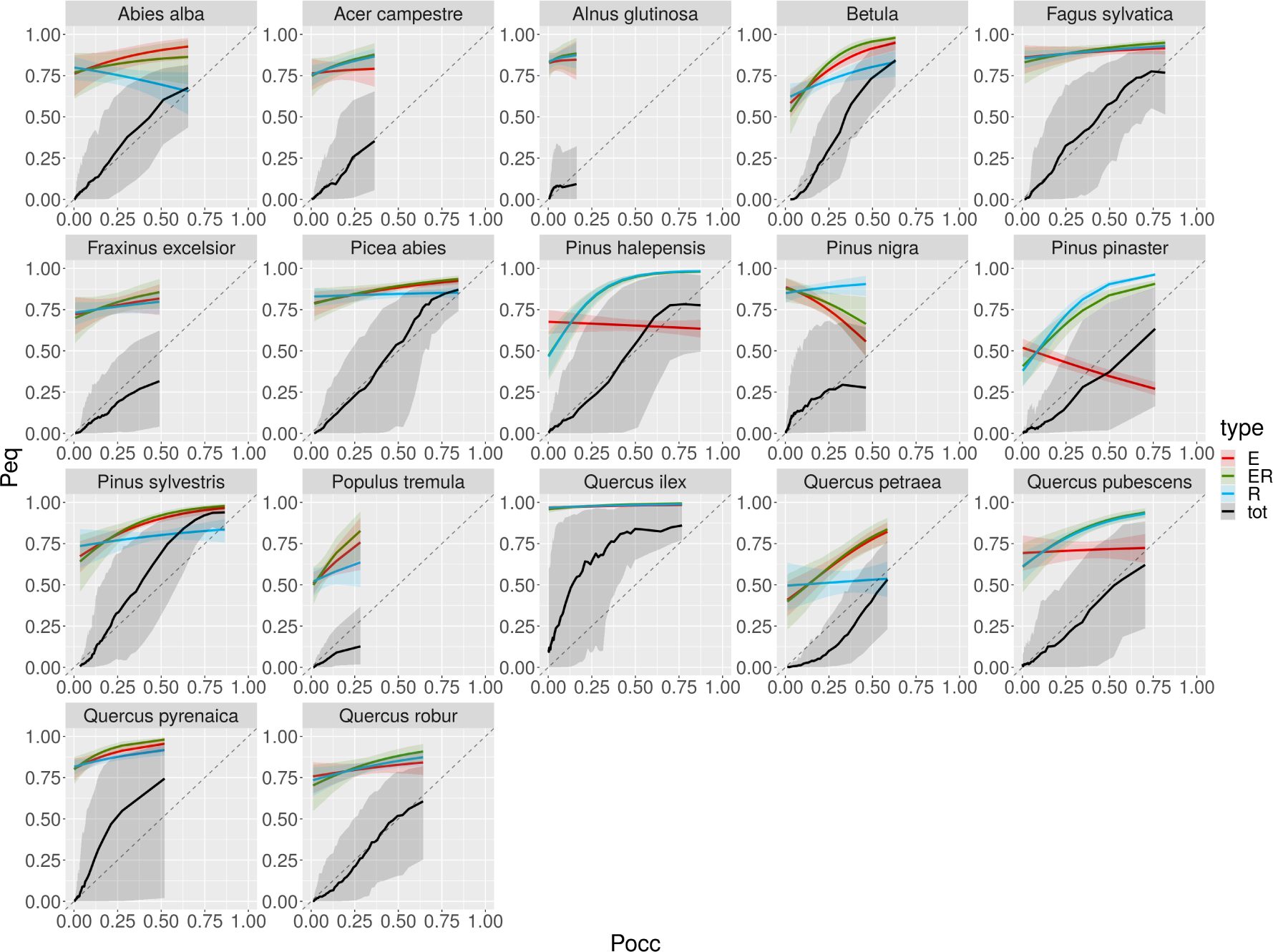
Equilibrium probabilities of presence (*P*_*eq*_) against SDM occurrence probability (*P*_*occ*_), calculated with open model with fixed seed source. Models either assume a seed source set to one and varying extinction (E in red), varying recruitment (R in blue), or both (ER in green) or extinction, recruitment, and seed source varying with *P*_*occ*_ (tot in back).

The second formulation of the equilibrium for a closed system leads also to a positive relationship between *P*_*eq*_ and *P*_*occ*_ (see green curves in Figure 5) with again the notable exception of *Pinus nigra*. In this case, an extinction probability higher than the recruitment probability would lead to a null value for *P*_*eq*_. Overall, we also found that *P*_*eq*_ overestimated *P*_*occ*_, and *P*_*eq*_ showed little variations along the *P*_*occ*_ gradient.

**Figure 5:**
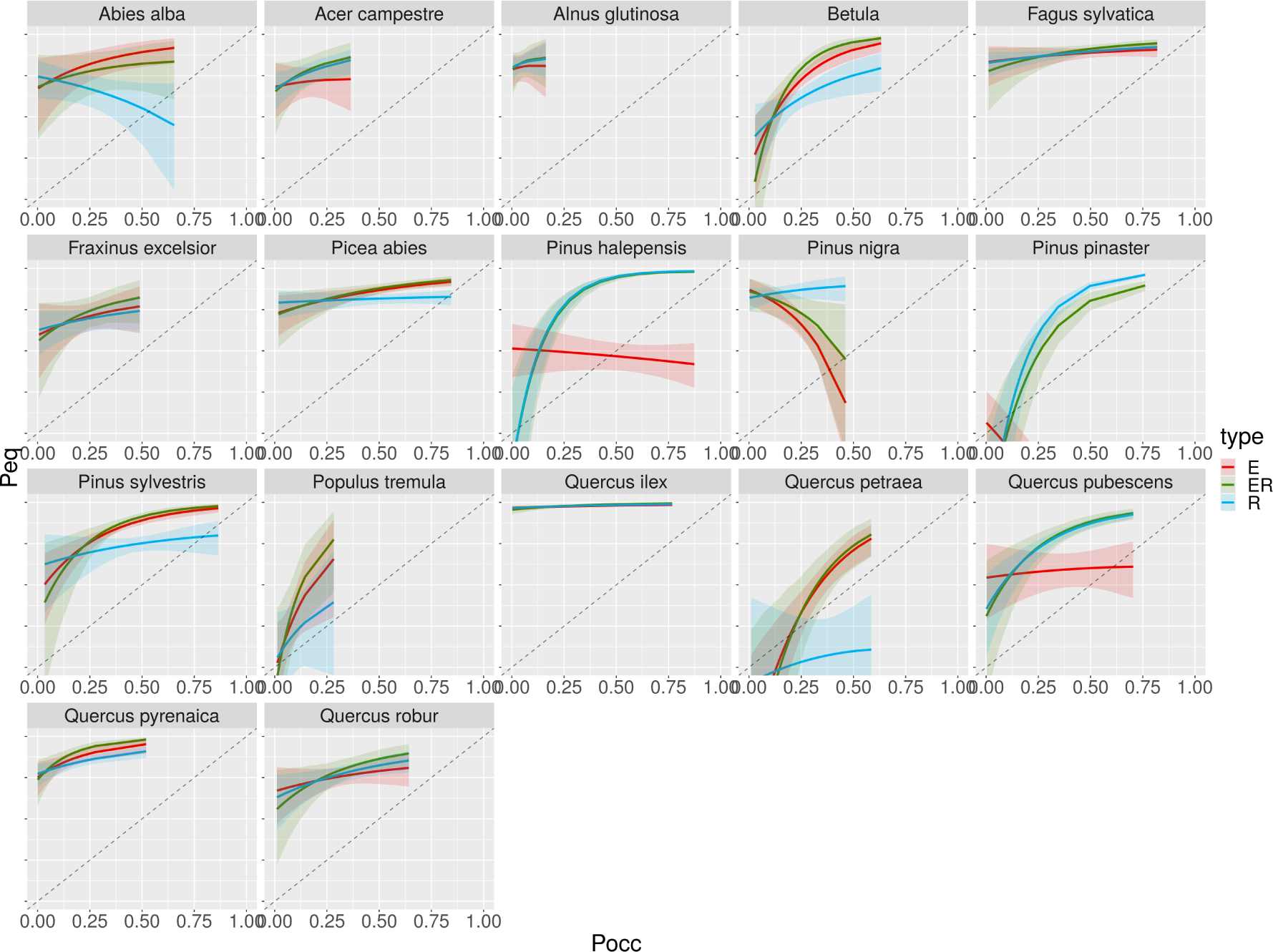
Equilibrium probability of presence (*P*_*eq*_) against SDM occurrence probability (*P*_*occ*_), calculated with closed formulation and varying extinction (E), varying recruitment (R), or both (ER).

## 4 Discussion

There is a long history of analyzing the drivers of population distribution, but surprisingly few studies have explored with field data the link between probability of presence and metapopulation processes such as extinction and colonization. Here, using data from national forest inventories, we explored the question at the scale of the European continent for 17 tree species. We found that for all species but *Fraxinus excelsior*, at least one of the processes, extinction or recruitment, is related to the SDM occurrence probability, but rarely both (only two species). When combining extinction and colonization, we also found that the probabilities of presence at equilibrium, derived from recruitment and extinction probabilities, were generally positively correlated with the observed occurrence probability (with the exception of *Pinus nigra*). However, at equilibrium, the metapopulation model generally overestimated the occurrence probability.

### 4.1 Variation of extinction and recruitment probability within species ranges

Holt and Keitt (2000) showed with theoretical models that there are different routes in metapopulation dynamics to range limits, via variations of colonization rates, variations of extinction rates, or variations of habitat availability. These three mechanisms are not mutually exclusive but can all be at play at the same time. Here, we directly explore the relative importance of the first two causes through the variations of recruitment and extinction. Our results reveal that variations within each species range were either through extinction or through recruitment but rarely both (only for *Quercus pyrenaica* and *Betula*). Generally, the magnitude of the response was stronger for recruitment than for extinction.

We found that for all species with a significant relationship between occurrence probability and extinction the relationship was negative, as expected by theory, except for *Pinus nigra* and *Pinus pinaster*. We found that for all species with a significant relationship between occurrence probability and recruitment, the relationship was positive as expected by theory. Thus only *Pinus pinaster* and *Pinus nigra* showed a significant response inverse to the theory for extinction, with an increase of the extinction probability with the increase of the occurrence probability (*Pinus halepensis* also showed a positive response but it was not significant). This opposite relationship for species belonging to the genus *Pinus* might be related to their intensive management and frequent plantation outside their native range (particularly in the case of *Pinus pinaster*). Current presence/absence data might be in that case biased to include location outside suitable habitats. The relationships between recruitment and occurrence probability were largely in agreement with the theoretical expectation as only *Abies alba* showed a negative but non-significant response.

There is no obvious explanation for why species respond through extinction or through recruitment. We found no clear explanation based on the species’ ecological strategies. There was no link between the slope of the response of recruitment or extinction and shade tolerance (using the shade tolerance index of Niinemets and Valladares (2006)) or key functional traits, see Supplementary Materials Section 7). Thuiller et al. (2014) proposed that shade tolerant species could show a closer relationship between population growth rate and SDM occurrence probability, as this link would be less blurred by competition, but they found, as do we, weak support for this hypothesis. We also verified that the SDM discrimination scores had no direct impact on mean slope estimations.

Relatively few other studies have explored with field observation of extinction and colonization if the metapopulation dynamics explain current species distribution (see for instance Talluto et al. (2017), García-Valdes et al. (2013), Araújo, Williams, and Fuller (2002)). These studies generally also supported the idea that metapopulation rates variations agreed with the species distribution, even if there was evidence of extinction debt and colonization credit at the species range (Talluto et al., 2017). Among these studies, the relative importance of extinction and colonization was not explicitly considered. The uncertainty of the estimation seems larger for the response of extinction to climate than for the response of colonization in Talluto et al. 2017, but it is not possible to compare the relative role of these two rates based on their results. Garcia-Valdes et al. 2015 found that climate had a stronger effect on extinction than colonization (whereas we found a stronger response of colonization). Overall there is a lack of studies exploring the relative magnitude of the variation of extinction and colonization within species ranges.

These previous studies on extinction-colonization probabilities of trees used patch occupancy models (Talluto et al., 2017; Purves et al., 2007; García-Valdes et al., 2013) with polynomial functions of climatic variables (such as temperature and aridity). A key difference with our model is that we did not use climatic variables directly but instead used the SDM occurrence probability as a descriptor of species environmental niches. Given the low number of colonization or extinction events in our data, using an SDM to summarize the species climatic niche might be more powerful than fitting complex multivariate responses to climatic variables. A similar approach has also been developed for birds in Britain by Araújo et al. (2002), and highlighted a negative relationship between local extinction probability and the occurrence probability.

More studies have focused on links between demographic rates and distribution. The links between species distribution and demographic rates (growth rate and carrying capacity by Thuiller et al. (2014), population growth rate, time to quasi-extinction, transient population dynamics by Csergő et al. (2017)) seem weaker than with metapopulation rates. This might indicate that links between population processes and species distribution are easier to capture with integrative metapopulation metrics than with detailed population-level metrics, which could be related to an issue of observation scale: colonization/extinction is a direct dynamical approach to presence/absence. In addition, upscaling individual demographic dynamics to presence/absence is not easy, due to non-linearity of demographic response to climate and temporal variability.

### 4.2 Implication for the probability of presence at equilibrium

Based on our analysis combining extinction and colonization to estimate the probability of presence at equilibrium, we found that in general the probability of presence at equilibrium was positively correlated to the occurrence probability estimated by the SDM, but with a strong overestimation. This was the case with both equilibrium formulations (open and closed). Thus, our estimates of extinction and colonization rates capture some drivers of the species environmental distribution but were not able to represent the current observed distribution. This agrees well with the previous patch occupancy model fitted to forest inventory data (Talluto et al., 2017; García-Valdes et al., 2013; García-Valdés et al., 2015) who also found that models were capturing part of the species range but with important deviations.

The two equilibrium formulations represent two extremes of the effect of seed dispersal. In the case of the closed formulation, the seed source outside the cell is not taken into account in the calculation. In that case, we see that our model would predict an increase in prevalence of most species, which can be related to the higher probability of recruitment compared to extinction across the occurrence gradient. In the open formulation, where seed input inside the cell is considered fixed and not affected by the metapopulation dynamics, we also found a strong overestimation of the occurrence probability from the SDMs. The most extreme case was *Quercus ilex* which showed strong overestimation of the mismatch between current and equilibrium probability of presence and very little variations with the open formulation. The only version of the model that did not strongly overestimate the probability of presence was the open formulation where the seed source varied according to the observed pattern within the species range. This latter formulation is strongly constrained the model by the current geographical distribution and provided little understanding of the mechanisms involved. This model might capture part of the last route to range limits proposed by Holt (Holt et al., 2005) because it explicitly took into account the proportion of forest/non-forest patches. However, a proper interpretation of these results would require to formally represent dispersal processes which was not possible in this study (see the discussion on this issue in the section on the limitations of patch occupancy models below).

The overestimation of the equilibrium probability of presence can arise because (1) the metapopulation processes are not in agreement with the current distribution and show some degree of non-equilibrium, or (2) our estimation of metapopulation dynamics and the colonization and extinction rate are not accurate enough. Below we discuss these two possible explanations.

### 4.3 Equilibrium *vs.* non-equilibrium of species distribution

If the distribution of a species was currently in equilibrium, we would expect a close match between the SDM and the probability of presence computed at equilibrium (due to either extinction, colonization, or both). It is important to note that equilibrium does not necessarily imply that both extinction and colonization processes are strongly related to SDM (see section 8 in Supplementary materials).

The fact that we are observing a positive correlation but not a perfect one to one relationship, however, does not rule out that there may be some degree of non-equilibrium between the metapopulation dynamics and the current distribution. The idea that each species is in current equilibrium with the environment has been criticized by Svenning and Skov (2004), based on the idea that most European tree species do not fully fill their potential ranges. This situation could be the result of a post-glacial migration lag as illustrated in Svenning, Normand, and Kageyama (2008). The lag would strongly affect *Abies alba*, the *Pinus* genus and the *Quercus* genus. This argument has however been partly contradicted by previous SDM results (Araújo & Pearson, 2005) and large dispersal rates found based on pollen records (Giesecke, Brewer, Finsinger, Leydet, & Bradshaw, 2017). Interestingly, we found a weak response of recruitment and extinction to SDM occurrence probability for *Abies alba*, a species with a recorded slow expansion rate (Giesecke et al., 2017). In Eastern North America, results from a SPOM model (Talluto et al., 2017) identified species out of equilibrium with climate at range margins. Their model formulation is close to our closed formulation (a species is present at equilibrium when colonization probability exceeds extinction probability). A SPOM developed by García-Valdes et al. (2013) in Spain also concluded on a non-equilibrium. Their simulations based on a model with constant climatic conditions lead to an increased fraction of occupied plots, but most species did not show range expansion.

It is important to stress that our analysis focused on testing whether metapopulation dynamics (colonization and extinction) were related to the SDM occurrence probability. Because we considered only a single gradient of occurrence probability we can not distinguish between changes at the southern or northern range and thus can not give an indication of directional range shift in comparison to patch occupancy model fitted with climatic variables.

### 4.4 Limitations of patch occupancy models

Several factors might have contributed to limit our ability to estimate the links between SDM and metapopulation dynamics and thus explain the mismatch between the equilibrium probability of presence and the SDM. First, the NFI data do not provide perfect informations on the absence/presence at the plot scale. With protocols that are based on concentric circular subplots for different size classes, we might miss the presence of trees larger than 10 cm DBH in one of the subplots. We partially corrected this issue, by accounting for the probability that a tree was below 10 cm at the first census with a growth model. But this approach is not perfect and the data set probably still contains colonization events that are not true colonization events but observation errors. Conversely, we might have wrongfully excluded some colonization events for trees with extreme growth. Using detailed recruitment data could improve our estimation, but they are not available for all NFI.

Another limitation is that our model did not explicitely consider dispersal. Different studies on patch occupancy models calibrated with NFI data have tried to formally include dispersal in the model (Purves et al., 2007; García-Valdés et al., 2015). García-Valdes et al. (2013) tried to infer the parameters of the dispersal kernel based on the Spanish forest inventory data. We considered that available knowledge on the potential seed source surrounding a plot is insufficient to draw mechanistic conclusions on seed dispersal. Field studies show that mean distances of seed dispersal are short for most tree species (Nathan et al., 2001; Bullock et al., 2017; Cain, Milligan, & Strand, 2000), therefore direct dispersal between plots should be restricted to extremely rare events (distance > 1 km) and the tail of the kernel distribution. It is thus very unlikely that these models were really estimating a dispersal kernel (as indicated by the very large mean dispersal distance inferred) but rather captured a degree of spatial auto-correlation in the species distribution and the recruitment process. Here, we use an estimate of local frequency which takes into account observed presence/absence and smoothing kernels as well as fine scale forest cover maps (building on the approach of Talluto et al. (2017)). We believe that if we want to include dispersal kernels in the model it is better to use external information on the shape and parameters of the dispersal kernel and have more accurate data on the seed source (see Schurr et al. (2007) or Schurr et al. (2012) for example).

Finally, our approach does not include biotic interactions and disturbances that might influence population extinction and recruitment probabilities (Case, Holt, McPeek, & Keitt, 2005; Svenning et al., 2014; Liang, Duveneck, Gustafson, Serra-Diaz, & Thompson, 2018). Trophic interactions in a broader sense may have a potentially large impact on recruitment estimation. For instance ungulate browsing may induce spatially varying limitations on recruitment for *Abies alba* (Kupferschmid, 2018), and ungulate preferences could lead to limitation of certain species (see e.g. stronger preference for *Pinus sylvestris* over *Pinus nigra* might reinforce *Pinus sylvestris* drought sensitivity, Herrero, Zamora, Castro, and Hódar (2012)). Given the small number of colonization and extinction events, a reliable estimate of tree species interactions with our data seems unrealistic.

## 5 Conclusion and perspectives

Several range dynamic models have already used SDMs to constrain metapopulation dynamics based on the assumption that occurrence probabilities derived from SDMs can be used as predictors of colonization and extinction rates (including range dynamics models and population viability analysis). Based on this assumption, SDM outputs are used either directly to define which grid cells are colonizable (see Engler and Guisan (2009)), or influence demographic information (Nenzén, Swab, Keith, & Araújo, 2012). Here we test this core assumption for 17 European tree species and found mixed support. At least one process, either colonization or extinction was related to the SDM, but generally not both and the match was far from perfect. We thus caution that models cannot simply assume that metapopulation dynamics is driven by SDM occurrence probability, but rather need to test which processes are affected and at which magnitude. Data driven patch occupancy models have the potential to go beyond criticized SDM correlative predictive approaches (Journé, Barnagaud, Bernard, Crochet, & Morin, 2019).

## 6 Acknowledgments

This work was funded under EU FP7 ERA-NET Sumforest 2016 through the call “Sustainable forests for the society of the future” (project REFORCE), with the ANR as national funding agency (grant ANR-16-SUMF-0002). The authors are grateful to Dr. Christian Wirth and Dr. Gerald Kändler for access to the harmonised inventory data. MAZ and PRB were supported grants DARE (RTI2018-096884-B-C32) and FUNDIVER (CGL2015-69186-C2-2-R); MICINN, Spain. The NFI data synthesis was conducted within the FunDivEUROPE project funded by the European Union’s Seventh Programme (FP7/2007–2013) under grant agreement No. 265171. We thank the MAGRAMA, the Johann Heinrich von Thunen-Institut, the Natural Resources Institute Finland (LUKE), the Swedish University of Agricultural Sciences, and the French Forest Inventory (IGN) for making NFI data available.

