## Supplementary Materials for "Colonization and extinction dynamics and their link to the distribution of European trees at continental scale"

July 8, 2020

#### 1 Country specific protocols

##### 1.1 Finnish National Forest Inventory

We used data from the eighth NFI (NFI8) of Finland sampled in the period 1985-1986 to 1995. For the inventory dataset we used information from plots surveyed in 1995. The sample plots are located in a systematic grid across the country of plot clusters in forested areas (Mäkipää & Heikkinen, 2003). In Southern Finland the grid is 16 km by 16 km square, with four plots in each cluster at 400 m intervals, while in Northern Finland the grid is a 24 km by 32 km rectangle with three plots per cluster, at 600 m intervals. These permanent plots were sampled using a variable radius technique with two concentric circular subplots of radius 5.64 m (i.e. 100 $m^2$) for trees with a diameter at breast height (DBH) below 10.5 cm and 9.77 m (i.e. 300 $m^2$) for trees with a DBH above 10.5 cm.

##### 1.2 French National Forest Inventory

The French NFI (IFN 2014) is based on a systematic 1 $km^2$ square grid covering the entire country. A forest is defined in the French NFI as a stand of more than 0.05 ha and wider than 20 m in which crowns from forest trees can reach 5 m and cover more than 10 percent of the area. The whole grid is measured in 5 years and the 5-year sample is divided into five systematic annual sub-samples. Trees are measured in three concentric plots, depending on their circumference at 1.3 m height. Trees with a circumference above 23.5 cm (corresponding to a DBH of 7.5 cm) are measured on a 6 m radius plot; trees with a circumference above 70.5 cm (DBH = 22.4 cm) are measured on a 9 m radius plot; trees with a circumference above 117.5 cm (DBH = 37.4 cm) are measured on a 15 m radius plot; trees with a circumference below 23.5 cm are not measured. For living trees, radial growth over the last five years is measured on short cores. See Bourdier et al. (2016) for details on the processing of the data to compute basal area growth at the plot scale. We used NFI data covering the 2007-2013 period. Growth measurements and evaluation of tree mortality during the past 5 years were used to reconstruct the stand structure 5 years before the measurements and then build two presence/absence datasets.

#### 1.3 German National Forest Inventory

We used information from the first and second German NFI. The German NFI uses a systematic grid of clusters, sampled during the periods 1986-1990 (undertaken in West Germany only) and 2001-2002. The size of the sample grid is 4 km by 4 km, however, it is reduced in some federal states to either 2.83 km by 2.83 km or 2 km by 2 km. Each cluster is a quadrangle of 150 m in length with a sample plot on each corner (Kandler, 2009). Trees with a DBH above 10 cm in the first inventory and above 7 cm in the second were selected by the angle-count method with a basal area factor (BAF) of 4 $m^2ha^{-1}$ if they are alive or recently dead.

#### 1.4 Spanish National Forest Inventory

We used information from the second and third Spanish NFI (surveyed during the periods 1986-1996 and 1997-2007, respectively). The Spanish NFI plots are located on a 1 $km^2$ grid over forested regions (Villaescusa & Díaz, 1998; Villanueva, 2004). Spanish NFI plots were sampled using a variable radius technique with four concentric circular subplots of radius 5 m for trees with DBH smaller than 12.4 cm, 10 m for those with DBH between 12.5 and 22.4 cm, 15 m for those with DBH between 22.5 and 42.4 cm and 25 m for those with DBH greater than 42.5 cm.

#### 1.5 Swedish National Forest Inventory

The permanent Swedish inventory uses a regular sampling grid and includes about 4,500 permanent tracts, each surveyed every five years. Plots in the first census were surveyed between 2003 and 2005 and plots in the second census were surveyed between 2008 and 2010. The tracts are rectangular and have different dimensions depending on the location within the country; each tract has between four and eight circular sample plots. All trees with a DBH above 10 cm are sampled in a 10 m radius.

| Country | Protocol method | Survey Dates | Plot radius/ minimum DBH | Plot radius/ minimum DBH | Plot radius/ minimum DBH | Plot radius/ minimum DBH |
|---|---|---|---|---|---|---|
| Spain | Variable radius | 1986/1996 1997/2007 | 5 m / 7.5 cm | 10 m / 12.5 cm | 15 m / 22.5 cm | 25 m / 42.4 cm |
| France | Variable radius | Yearly 20052011 | 6 m / 7.5 cm | 9 m / 22.5 cm | 15 m / 37.5 cm | |
| Sweden | Variable radius | 2005/2010 2008/2010 | 10 m / 10 cm | | | |
| Finland | Variable radius | 1985/1986 1995 | 5.64 / <10.5 cm | 9.8 m >10.5 | | |
| Germany | Angle count 4 $m^2 ha^{-1}$ | 1986/1990 2001/2002 | | | | |

### 2 Growth Model

In order to correct for potential recruitment error, we used a species-specific growth model to assess the probability distribution of the DBH of a recruited tree. If the probability that the former DBH was below 10 cm exceeds 0.5, we consider the new tree as a recruitment. The growth model was built using a glm with two climatic covariates (a water aridity index and the sum of growing degree days) extracted from E-OBS, a high resolution (1 $km^2$) downscaled climate data-set (Moreno & Hasenauer, 2015) and as a function of tree DBH. See Kunstler et al. (2019) for more detailed information on the growth model. A plot random effect is also considered in the model. We verified that model residuals did not exhibit bias to ensure we do not have climatic bias in colonization observation corrections, (see for example figure 1). Table 1 resumes each species-specific evaluation via the marginal pseudo R squared.

| Species | Marginal R2 |
|---|---|
| *Pinus sylvestris* | 0.14 |
| *Picea abies* | 0.30 |
| *Fagus sylvatica* | 0.33 |
| *Quercus robur* | 0.19 |
| *Quercus petraea* | 0.30 |
| *Pinus pinaster* | 0.23 |
| *Quercus ilex* | 0.11 |
| *Pinus nigra* | 0.19 |
| *Abies alba* | 0.25 |
| *Pinus halepensis* | 0.21 |
| *Quercus pubescens* | 0.19 |
| *Betula* | 0.20 |
| *Fraxinus excelsior* | 0.26 |
| *Quercus pyrenaica* | 0.19 |
| *Alnus glutinosa* | 0.18 |
| *Populus tremula* | 0.22 |
| *Acer campestre* | 0.23 |

Table 1: Marginal R-squared value (accounting for fixed effect) for each species.

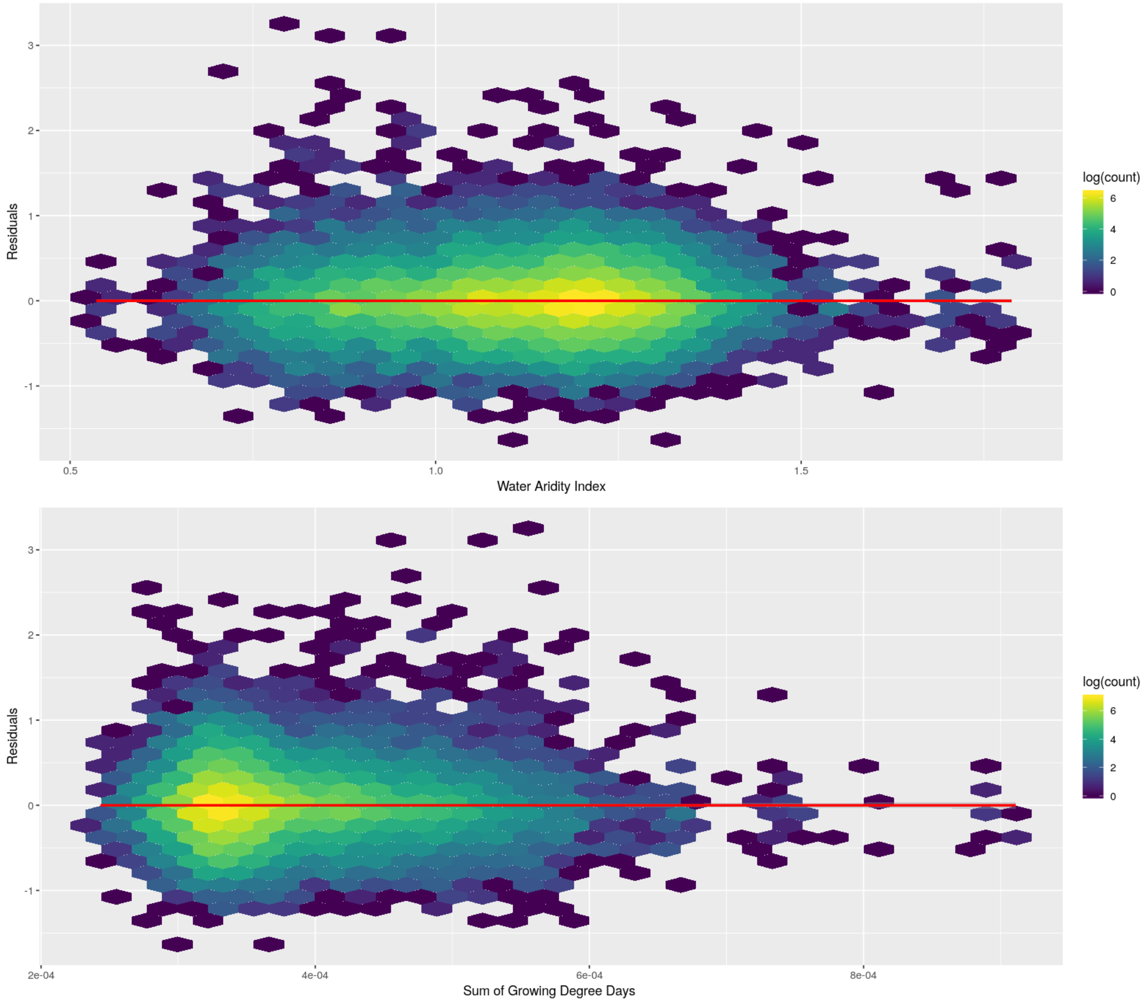


Figure 1: Growth model residuals as a function of the two climatic variables for *Quercus pubescens*

#### 3 SDM evaluation

For each species, probability of presence is fitted using an ensemble model using BIOMOD2 R package (Thuiller et al., 2016). Each model (Generalized Linear Models, Generalised Additive Models, Generalized Boosted Models and Random Forests) are evaluated using two discrimination scores: Area Under the Curve (AUC) and True Skill Statistic (TSS). Each model was fitted five times using 70 % of available data and evaluated on the 30 % remaining data. Results are summarised in figure 2. Both AUC and TSS statistics evaluate the discrimination potential of a model. AUC ranges from 0 to 1, 0.5 being the score of a null model; TSS ranges from -1 to 1, 0 being the score of a null model.

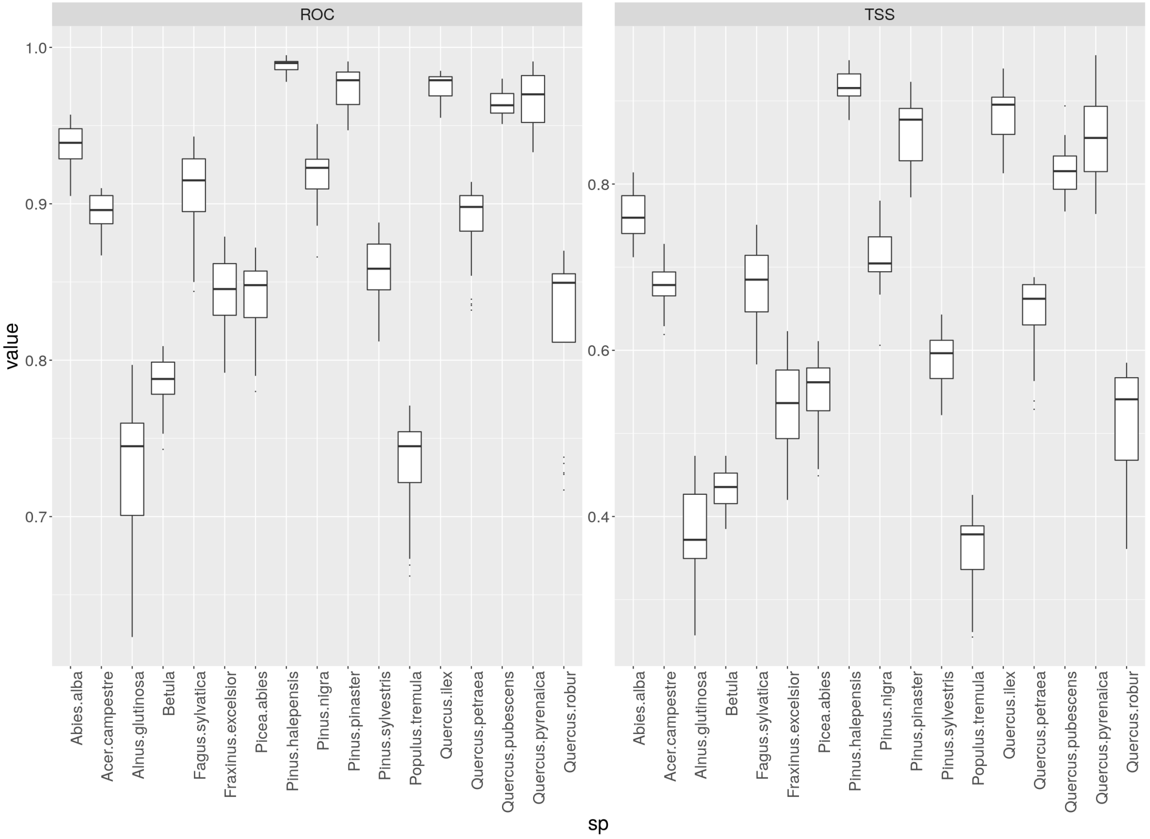


Figure 2: Discrimination scores (Area Under the Curve on the left, True Skill Statistics on the right) of SDM per species. Boxplots summarise the scores from different model formulation (GLM, GAM, GBM and RFF) and samplings.

SDM projections were done by averaging the outputs of the best models ($TSS > 0.4$).

### 4 Links between SDM performance and recruitment and extinction probability slopes

We found no significant relationship between SDM performance (Area Under the Curve) and the mean slopes estimated by our model for the response of extinction and colonization to SDM probability of occurrence (top figures in 3). However SDM performances had an impact on posterior distribution, with a larger standard deviation of the posterior distribution when the SDM score is low (bottom figures in 3).

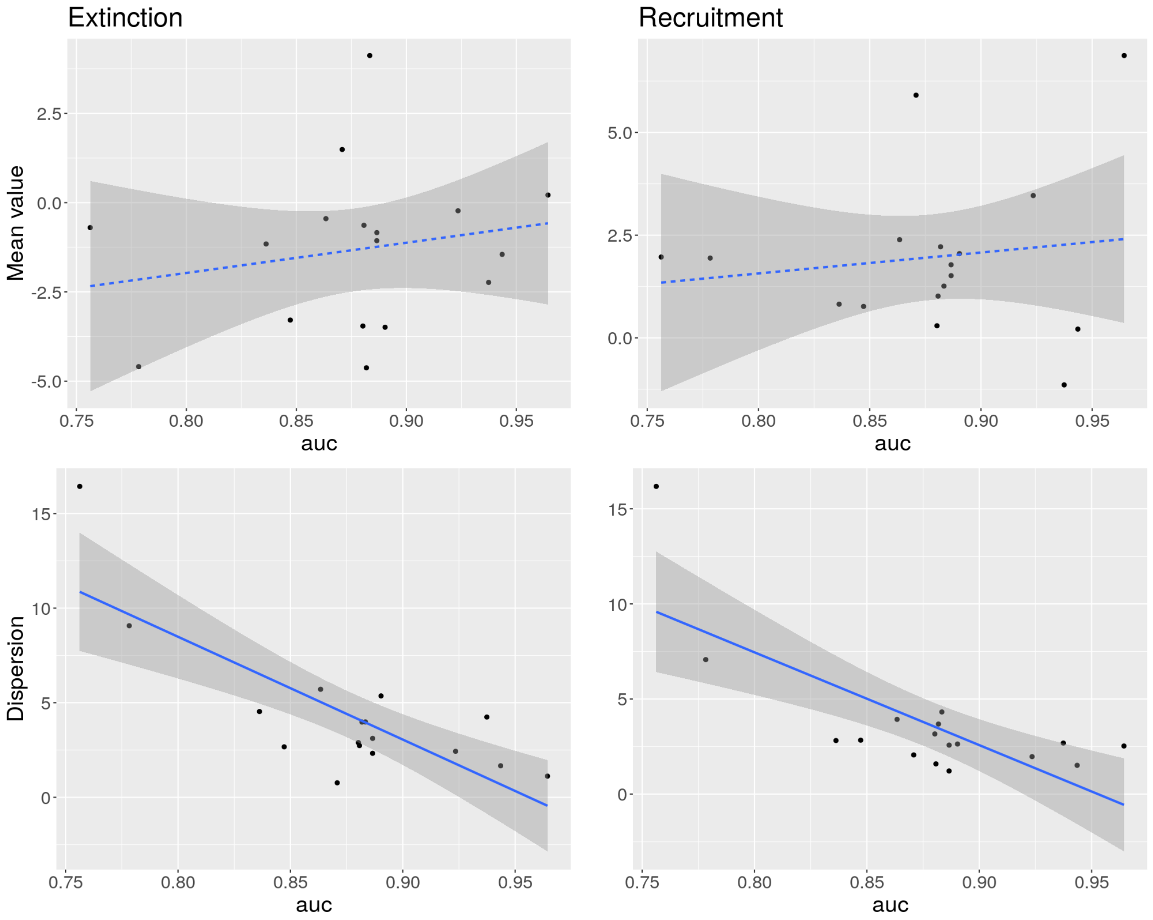


Figure 3: Extinction (left) and recruitment (right) relationship between SDM performance score (AUC) and (top) the mean posterior distribution of the slope and (bottom) the standard deviation of the posterior distribution slopes. Dotted lines mean the relationship is not significant at 5 %.

### 5 Equilibrium probability variations with the probability of occupancy

We evaluated how different combinations of parameters values for the extinction and colonization probability (including intercept and slope parameters) affect the shape of the relationship $P_{eq}$ vs. $P_{occ}$. The figure below present this using the open equilibrium formulation. The figure clearly shows that a range of shapes is possible.

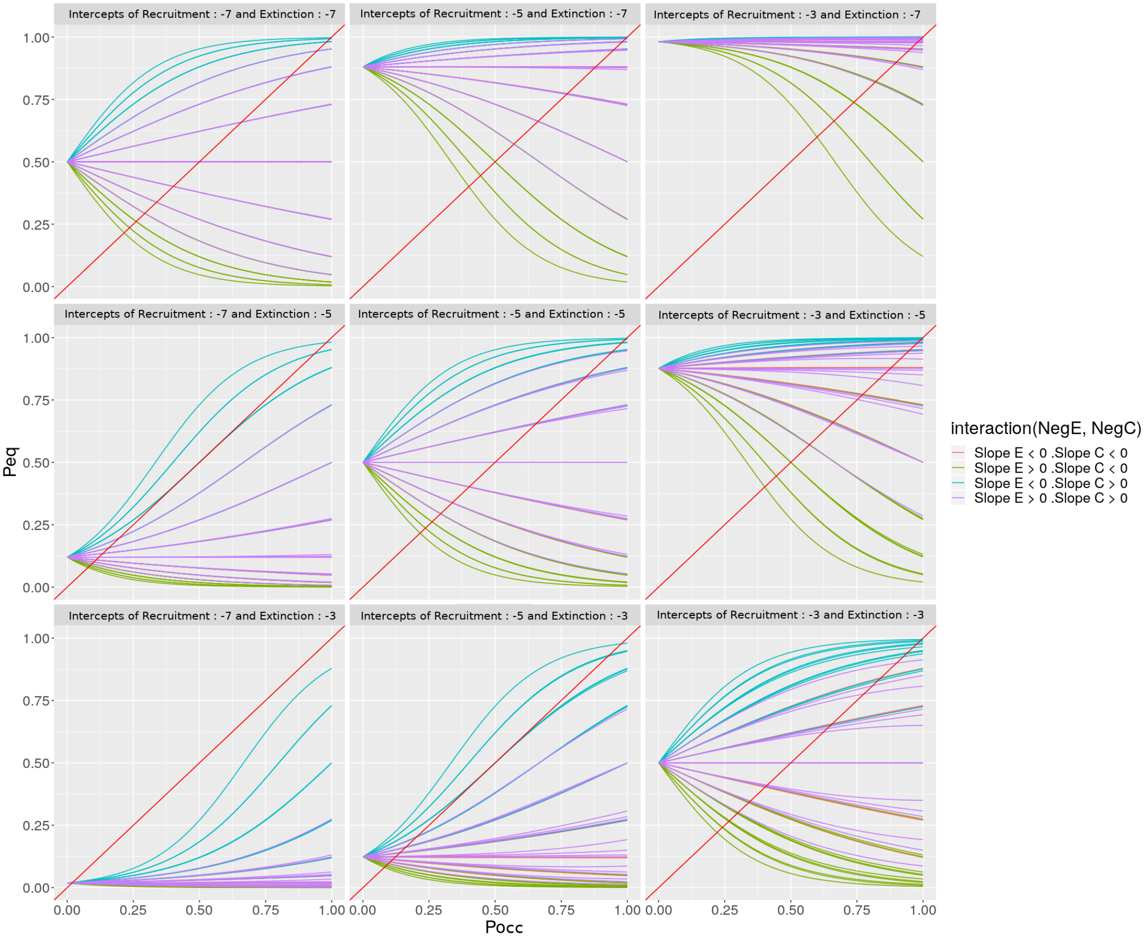


Figure 4: Peq as a function of Pocc with varying extinction/recruitment model parameters. Intercepts vary between panels, and within each panel, slope parameters can take integers values from -3 to +3. Most species lie in the case of negative extinction slope and positive recruitment slope (blue lines), with a higher recruitment intercept than extinction intercept (upper right panels)

### 6 Rare events impact

To test the potential impact of the low proportions of colonization/extinction events, we performed some tests using all positive events (1s) and sampled a variable fraction of negative events (0s). For each fraction (plots retained vary from $\frac{2}{8}$ to $\frac{7}{8}$ of total observations), we performed 100 samples of 0s and computed the slope parameter by maximizing the likelihood function. We then took the mean of these 100 samplings, for recruitment probabilities (Figure 5) and extinction probabilities (Figure 6). These two figures show that the effect of using a smaller sample of 0s is relatively weak with effect only with a very low number of 0s for recruitment. Overall this analysis shows that our results are robust to the number of 0s. Our slope estimates are thus not strongly influenced by the relative small proportion of 1s.

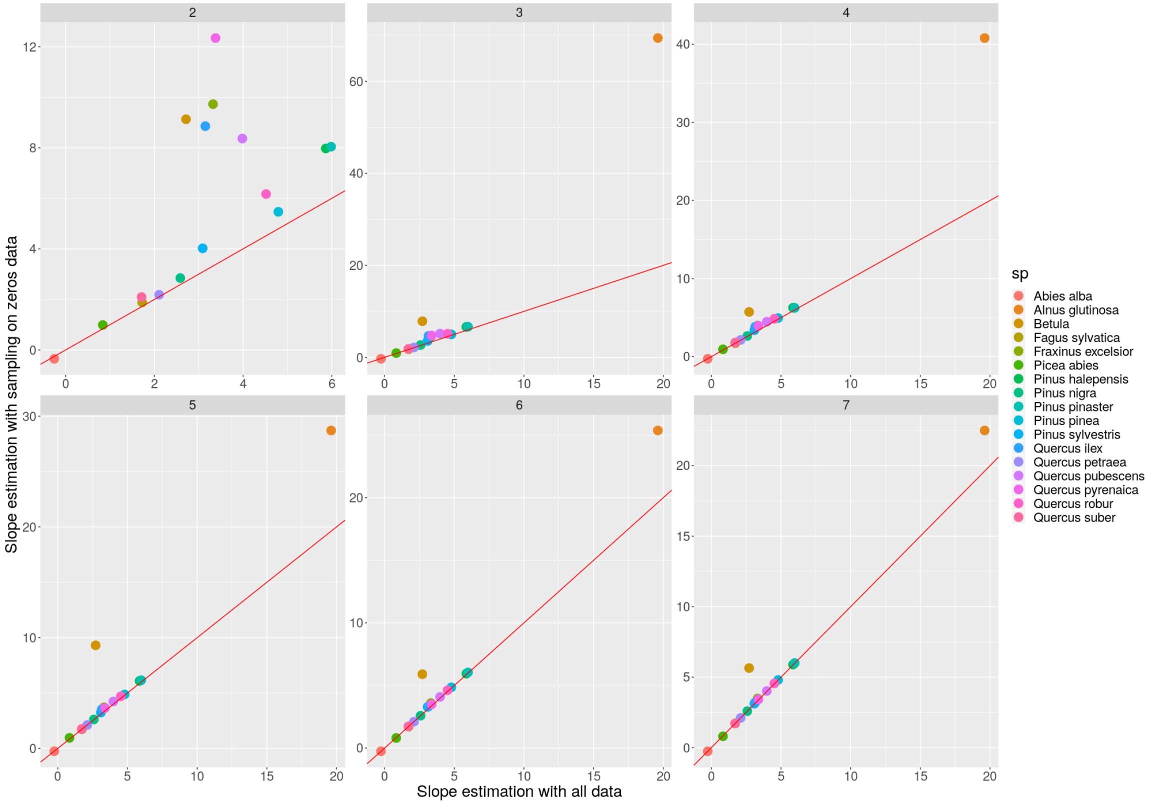


Figure 5: Slope of recruitment rates estimated with the full data set (x-axis) compared with slopes estimated (y-axis) with varying proportion of 0s (from 2/8 upper left to 7/8 bottom right).

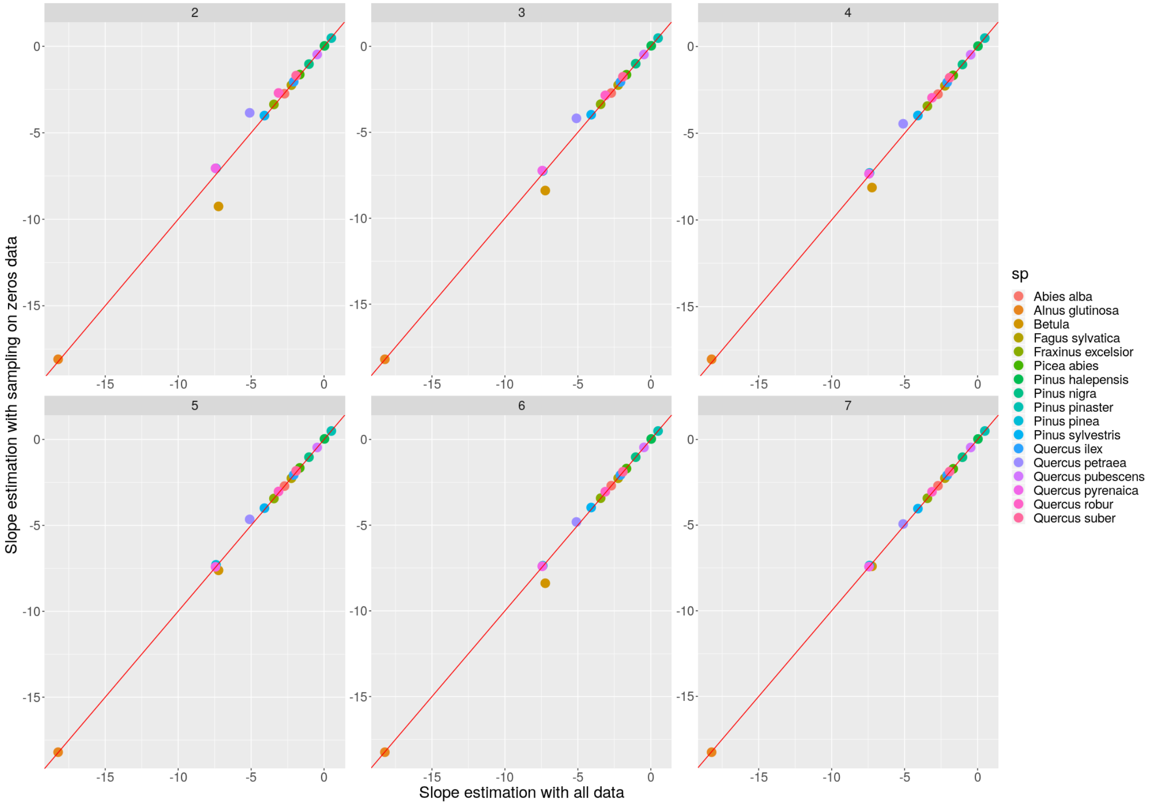


Figure 6: Slope of extinction rates estimated with the full data set (x-axis) compared with slopes estimated (y-axis) with varying proportion of 0s (from 2/8 upper left to 7/8 bottom right).

### 7 Metapopulation rates and traits

We tested whether potential species ecological strategy could explain the differences observed in extinction and recruitment relationship with $P_{occ}$. We found no clear relationship between model slopes and shade tolerance index (figure 7). In addition we conducted the same test with four key functional traits (figure 8) extracted from open databases (Wright et al., 2004; Chave et al., 2009; Choat et al., 2012; Maire et al., 2015): the specific leaf area and the leaf nitrogen per mass for leaf related traits, wood density and the xylem vulnerability to embolism measured by the water potential leading to 50% loss of xylem conductivity. All effects were negligible especially after a Bonferroni correction to account for multiple comparisons.

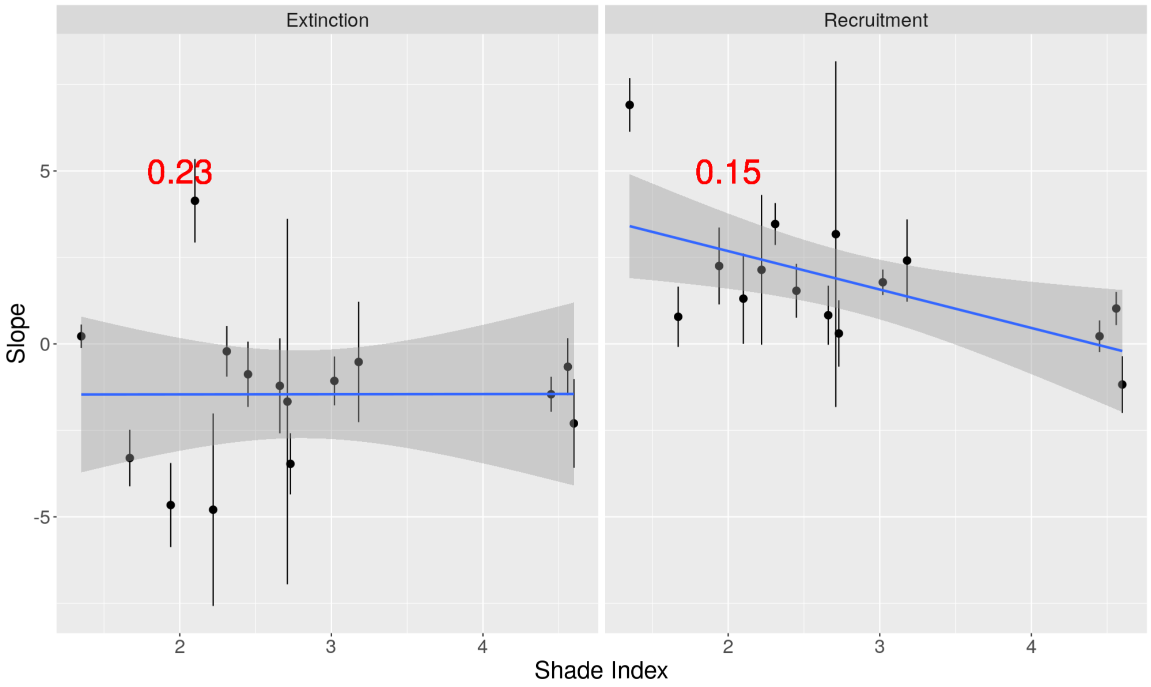


Figure 7: Recruitment slopes (left) and extinction slopes (right) against shade tolerance index from Niinemets and Valladares (2006). Number in reds are p-values associated with the shade tolerance index slope.

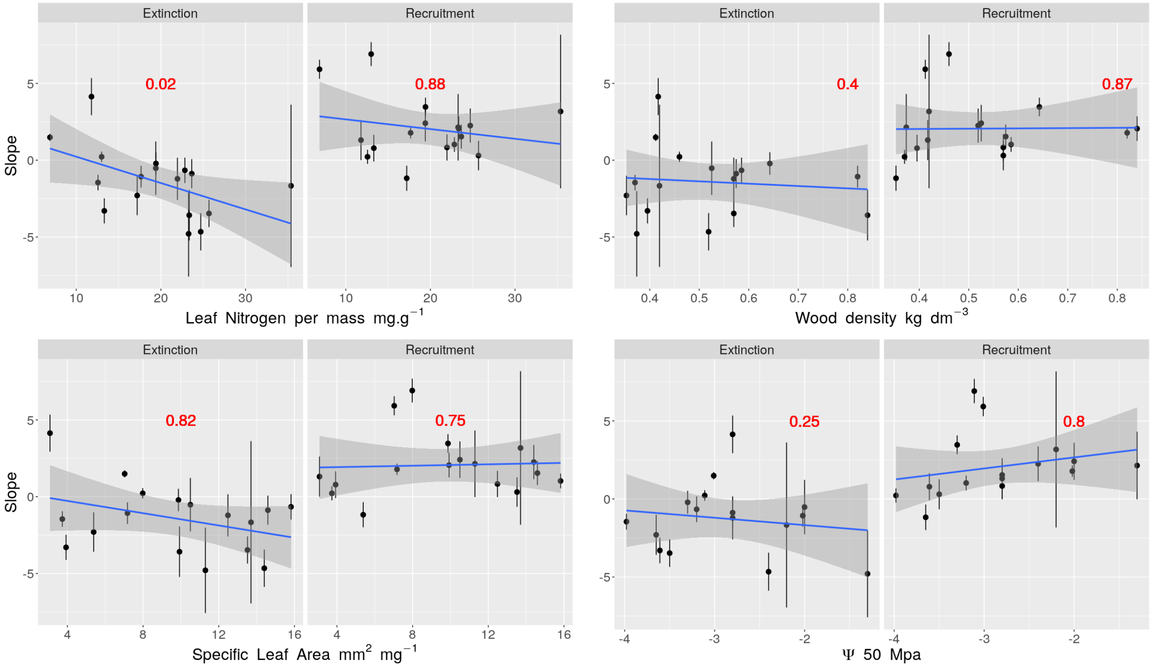


Figure 8: Recruitment slopes (left) and extinction slopes (right) against four traits : Leaf nitrogen per mass (upper left), Stem specific density (upper right), Specific leaf area (bottom left) and Ψ50 (bottom right). Number in reds are p-values associated with the slope.

### 8 On the links between extinction/colonization probabilities with equilibrium probability of presence

Here we explore whether metapopulation extinction and colonization probabilities, E and C respectively, are necessarily correlated with probability of presence $P_{eq}$ for a metapopulation without external seed source, when we assume that the population is at equilibrium. We will show that, even based on this equilibrium assumption, one of C or E can be completely uncorrelated with $P_{eq}$ , and the other probability exhibited limited correlation ($R^2 < 0.4$). We stress that for our analysis of European tree species, we do not know whether their metapopulations are close to equilibrium or not (Svenning & Skov, 2004) so the links are probably even less constrained. The equilibrium condition for a metapopulation without external seed source is given as: $P_{eq} = \frac{C}{C+E}$. This can be rearranged to yield : $logit(P_{eq}) = log(\frac{C}{C+E})$.

We model $P_{eq}$ along a single environmental gradient x ranging from -2 to 2, with $logit(P_{eq})$ as a quadratic function of x, which can be seen as a simple case for a species distribution model.

We could model $log(E)$ and $log(C)$ also as linear functions of the environment. However, this might lead to values of E and C larger than 1. We consider the case where the extinction rate linearly increases along the environmental gradient (i.e. increasing extinction at the warm edge). Therefore, we model the logarithms of the rates e and c as linear functions of the environment. Rates and probabilities are related by $E = 1 - exp(-e\Delta t)$, where $\Delta t$ is the time step. s We calculate the probability E, assuming that the time step is small relative to the units in which e is expressed (e.g. if e was in 1/century, and the time step was one decade, then $\Delta t$ would be 0.1)

We can now directly calculate the parameters of the colonization rate, assuming that we can approximate $log(\frac{C}{C+E})$ well with $log(\frac{c}{c+e})$ - we confirm that assumption visually below.

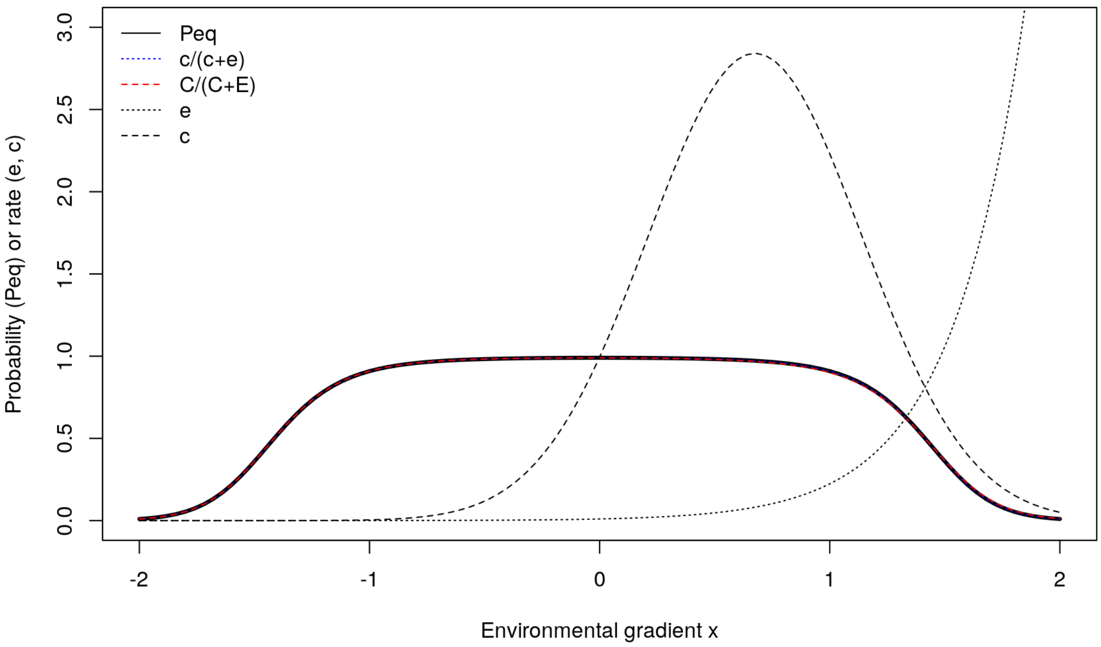


Figure 9: Occurrence probability $P_{eq}$ , extinction and colonization rates e and c along the environmental gradient x.

The estimates of $P_{eq}$ based on rates (c and e) and probabilities (C and E) agree well. The extinction rate is visually uncorrelated with the occurrence probability; the maximum of the colonization rate is shifted towards higher environmental values relative to the maximum of the occurrence probability, to compensate for the increasing extinction rates at the warm edge. Finally, we estimate the R-squared of models of $logit(E)$ and $logit(C)$ as linear functions of $P_{eq}$ (assuming a Gaussian error) and found a R-squared of 0 between $logit(E)$ and $P_{eq}$, and 0.36 between $logit(C)$ and $P_{eq}$.
